# Intravital imaging of islet Ca^2+^ dynamics reveals enhanced β cell connectivity after bariatric surgery in mice

**DOI:** 10.1101/2020.05.05.078725

**Authors:** Elina Akalestou, Kinga Suba, Livia Lopez-Noriega, Eleni Georgiadou, Pauline Chabosseau, Isabelle Leclerc, Victoria Salem, Guy A. Rutter

**Author notes:** **Corresponding Authors:** Guy A. Rutter, Room 327, ICTEM Building, Imperial College London, Hammersmith Hospital Campus, Du Cane Road, W12 0NN London, United Kingdom, Victoria Salem, 6^th^ Floor Commonwealth Building, Imperial College London, Hammersmith Hospital Campus, Du Cane Road, W12 0NN London, United Kingdom. **Author Contributions:** E.A., K.S., V.S. and L.L.N. undertook the mouse studies. V.S. and G.A.R. designed and supervised the study. E.A. undertook all data analyses. E.G. developed connectivity scripts and P.C. contributed to connectivity analysis. I.L assisted with mouse studies. E.A. and G.A.R. wrote the manuscript with contributions from all authors.

## Abstract

Bariatric surgery improves both insulin sensitivity and secretion in type 2 diabetes. However, these changes are difficult to monitor directly and independently. In particular, the degree and the time course over which surgery impacts β cell function, versus mass, have been difficult to establish. In this study, we investigated the effect of bariatric surgery on β cell function *in vivo* by imaging Ca^2+^ dynamics prospectively and at the single cell level in islets engrafted into the anterior eye chamber. Islets expressing GCaMP6f selectively in the β cell were transplanted into obese male hyperglycaemic mice that were then subjected to either vertical sleeve gastrectomy (VSG) or sham surgery. Imaged *in vivo* in the eye, VSG improved coordinated Ca^2+^ activity, with 90% of islets observed exhibiting enhanced Ca^2+^ wave activity ten weeks post-surgery, while islet wave activity in sham animals fell to zero discernible coordinated islet Ca^2+^ activity at the same time point. Correspondingly, VSG mice displayed significantly improved glucose tolerance and insulin secretion. Circulating fasting levels of GLP-1 were also increased after surgery, potentially contributing to improved β cell performance. We thus demonstrate that bariatric surgery leads to time-dependent increases in individual β cell function and intra-islet connectivity, together driving increased insulin secretion and diabetes remission, in a weight-loss independent fashion.

**Significance Statement:** Used widely to treat obesity, bariatric surgery also relieves the symptoms of type 2 diabetes. The mechanisms involved in diabetes remission are still contested, with increased insulin sensitivity and elevated insulin secretion from pancreatic β cells both implicated. Whilst the speed of remission – usually within a few days – argues for improvements in β cell function rather than increases in mass, a direct demonstration of changes at the level of individual β cells or islets has been elusive. Here, we combine vertical sleeve gastrectomy with intravital imaging of islets engrafted into the mouse anterior eye chamber to reveal that surgery causes a time-dependent improvement in glucose-induced Ca^2+^ dynamics and β cell - β cell connectivity, both of which likely underlie increased insulin release.

## Introduction

An estimated 30 million individuals in the US (9.4 % of the population) have diabetes (1), with ∼90% of cases thought to be Type 2 Diabetes (T2D), while in the United Kingdom it is predicted that by 2025 more than five million people will be diagnosed with the disease (2). In response to this epidemic, an abundance of pharmacological, dietary, exercise and behavioural interventions have been deployed but often focus on T2D management rather than long-term disease resolution (3, 4). Several clinical trials have now reported that bariatric surgery, a group of gastrointestinal procedures originally developed to aid weight loss, improves long-term glycaemia more effectively than caloric restriction or medical intervention (5-7).

Numerous studies (8-12) have attempted to unravel the mechanisms through which blood glucose control is improved post-operatively. One hypothesis to explain post-bariatric T2D remission is that it results from the increased release of incretins from the gut, such as the gastrointestinal insulin-stimulating hormone Glucagon-like Peptide 1 (GLP-1), as upregulated postprandial levels have been reported following bariatric surgery (13-15). Preclinical and clinical data have shown that bariatric surgery improves both hepatic and peripheral insulin sensitivity, as well as increases in insulin secretion (16-19). However, the exact mechanisms through which surgery impacts the β cell, including the identity of all the extra-pancreatic signals involved, and the relative importance of changes in β cell function and mass, have remained elusive. Nonetheless, the rapid (hours-days) reversal of diabetes in human subjects treated with bariatric surgery (20, 21) has provided powerful evidence that an improvement of β cell function plays an important, and possibly the dominant, role in increasing pancreatic insulin output.

A critical limitation in investigating β cell function in living humans or preclinical models is that, in the absence of robust *in vivo* imaging technologies (22), function must rely mainly on indirect measurements of circulating insulin or C-peptide. These approaches preclude any quantitation of changes over time, a detailed examination of function at the level of single β cells, or the connections between them. The latter has become an important issue since we (23) and others (24) have reported that weaker intercellular connections, and the loss of highly connected cells, that can often initiate Ca^2+^ waves (sometimes referred to as “hubs”), underlie the loss of insulin secretion observed in response to challenges associated with diabetes gluco(lipo)toxicity, low inflammation level, etc. (23, 25, 26). However, untangling these functional changes from alterations in β cell mass *in vivo* is challenging, since the latter can only reliably be determined post-mortem via pancreatic biopsies, and thus at a single time point.

In an effort to overcome these limitations, the present study aimed to investigate the effect of Vertical Sleeve Gastrectomy (VSG) on pancreatic β cell function in mice, by transplanting “reporter” islets in the anterior chamber of the eye. This approach was established by Berggren and colleagues (27) and has recently been developed by ourselves (26) to assess coordinated islet behaviour *in vivo*. Importantly, this technique has allowed us to image Ca^2+^ dynamics recursively, in the same islet, over time and with near single cell resolution, following surgery. We show that VSG increases β cell Ca^2+^ dynamics within eight weeks post-surgery when compared to pre-operative baseline and a sham operated group. Moreover, we demonstrate that VSG increases the number and strength of β to β cell connections at ten weeks after surgery. These changes were associated with increased fasting levels of GLP-1, suggesting that enhanced incretin production may contribute to postoperative improvements in β cell performance.

## Results

### Vertical Sleeve Gastrectomy improves glucose tolerance

Our experimental protocol is summarized in Figure 1A. In brief, mice were placed on a high fat high sucrose diet (HFHSD), at eight weeks of age, eight weeks before sham or vertical sleeve gastrectomy (VSG) surgery (week 0). This protocol led to fasting hyperglycaemia, indicative of β cell decompensation and defective insulin secretion, as expected (28). Ins1Cre:GCaMPf^fl/fl^ islets were isolated from donor mice and transplanted at week (−4). Baseline islet Ca^2+^ dynamics were imaged at week (−1).

**Figure 1:**
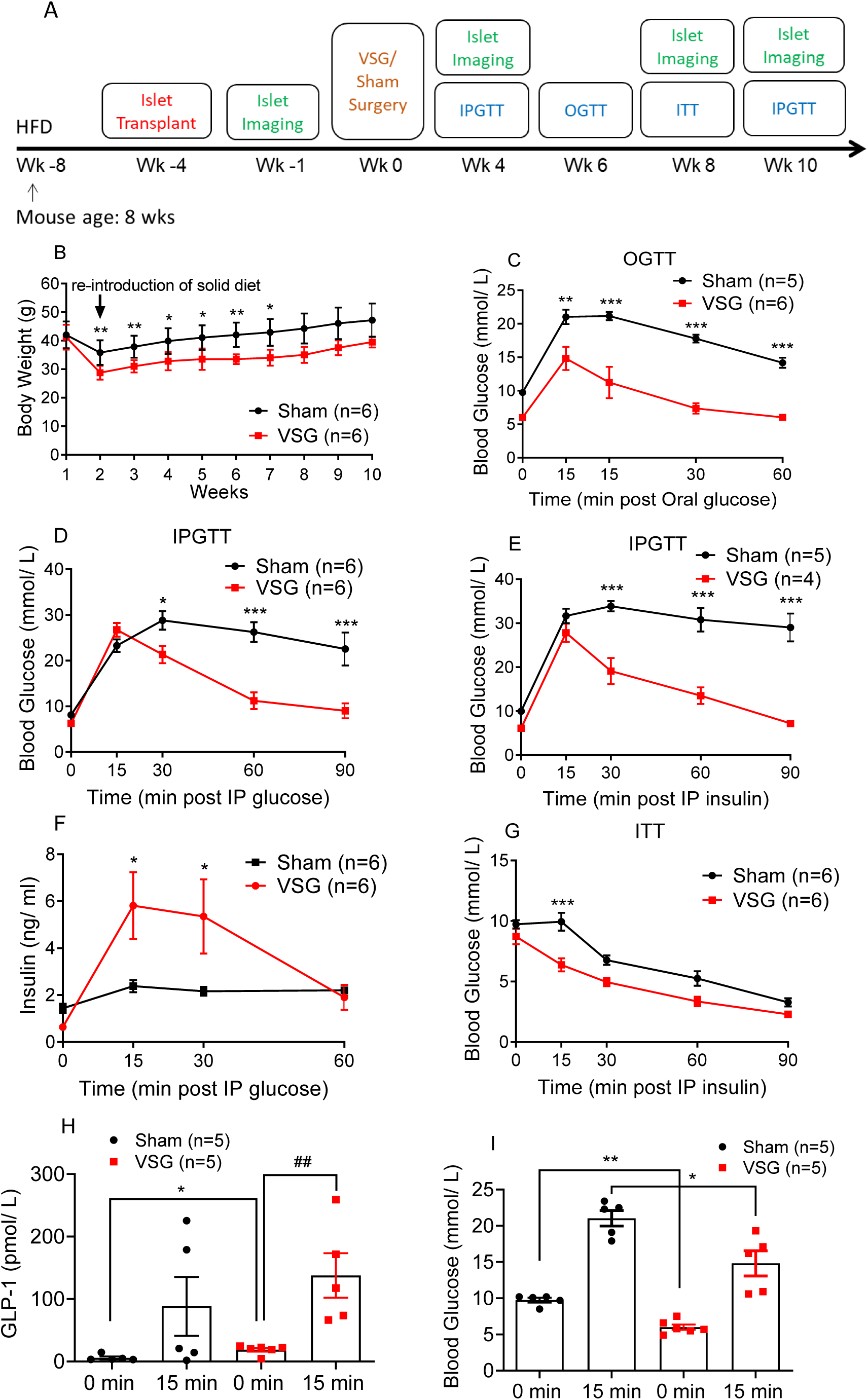
VSG improves glucose and insulin tolerance in HFHSD mice. (A) Timeline of procedures. (B) Body weight monitoring following VSG (n=6 animals) or sham surgery (n=6). (C) Glucose was administered via oral gavage (3 g/kg) after mice were fasted overnight and blood glucose levels measured at 0, 15, 30, 60 and 90 min. post gavage, six weeks after surgery, n=5-6 mice/group. (D) Glucose was administered via intraperitoneal injection (3 g/kg) after mice were fasted overnight and blood glucose levels measured at 0, 15, 30, 60 and 90 min. post injection, four weeks after surgery, n = 6 mice/group. (E) Glucose was administered via intraperitoneal injection (3 g/kg) after mice were fasted overnight and blood glucose levels measured at 0, 15, 30, 60 and 90 min. post injection, 10 weeks after surgery, n = 4-5 mice/group. (F) Corresponding insulin secretion levels measured on plasma samples obtained during the IPGTT performed in D (n=6). (G) Insulin was administered via intraperitoneal injection (1.5UI/kg) after mice were fasted for 5h and blood glucose levels measured at 0, 15, 30, 60 and 90 min. post injection, 7-8 weeks after surgery, n = 6 mice/group. (H) Corresponding GLP-1 secretion levels measured on plasma samples obtained during the OGTT performed in C (n=5). (I) Corresponding glucose levels for 0, 15 min obtained during the OGTT performed in C (n=5). ## P<0.01 VSG 0 vs. VSG 15 min, *P<0.05, **P<0.01,***P<0.001 VSG vs. Sham, following Student t-test or 2-way ANOVA. Data are expressed as means ± SEM.

VSG-treated mice experienced a larger decrease in body weight versus sham-operated animals, that was statistically significant until week 8 (week 7 av. Sham 42.9 ± 4.3g, av. VSG 34 ± 2.4g, p>0.05), (Fig. 1B). Vertical sleeve gastrectomy significantly increased the glucose clearance rate (p<0.01 at 15, 30, 60 and 90 min.) as assessed by oral glucose tolerance test (OGTT) at post-operative week 8 (Fig. 1C) and intraperitoneal glucose tolerance test (IPGTT) four and ten weeks post operatively (p<0.01 at min. 30, 60, 90 min, Fig. 1D, 1E respectively). Strikingly, in all tolerance tests performed on VSG-treated mice, glucose peaked at 15 min. post glucose injection (3g/kg) and dropped to baseline levels within 60 min. by week eight, and near baseline levels at week ten. In contrast, in sham-operated mice, glucose peaked at 30 min. and did not fully recover within the first 2 h of measurement.

### Vertical Sleeve Gastrectomy improves insulin secretion and sensitivity but does not increase β cell mass

In order to understand the marked increase in the rate of glucose clearance in mice that had undergone VSG, we measured insulin secretion *in vivo* as a response to an IP glucose load (3g/kg). Insulin secretion was increased significantly in VSG versus sham mice as early as four weeks post operatively (Fig. 1F, with the observed peak at 15 min. almost three-fold higher compared to sham mice (p<0.05). VSG mice were also significantly more insulin sensitive when compared to sham mice, as assessed by intraperitoneal insulin tolerance test (ITT, 1.5U/kg) (p<0.01, Fig. 1 G). However, pancreatic β cell mass was not increased in the VSG group relative to sham controls (Supp. Fig. 1A, 2). Notably, the ratio of α to β cell mass was significantly higher in the VSG group, yet α cell mass was not significantly increased (Supp Fig. 1B, C, 2).

### Vertical Sleeve Gastrectomy enhances GLP-1 secretion

To assess whether enhanced incretin release may contribute to the euglycemic effect of VSG we observed during IPGTT and OGTT, we measured plasma GLP-1 levels during fasting and 15 min. following an orally administered glucose load (3g/kg) (Fig. 1C). Fasting GLP-1 was significantly higher in the VSG group, when compared to sham (Fig. 1H). Moreover, whilst glucose failed to increase GLP-1 levels significantly in the sham group, a highly significant increase in response to glucose gavage was observed in VSG-treated animals (Fig. 1H). Significantly lower glucose levels were apparent in VSG-treated versus sham-treated mice, both fasting and following glucose gavage (Fig. 1I).

### β cell Ca^2+^ dynamics are enhanced following Vertical Sleeve Gastrectomy

In order to explore changes in β cell function after surgery, we monitored intracellular Ca^2+^ changes prospectively and in the same islets by confocal imaging of the anterior eye chamber (26, 27). Ca^2+^ increases, measured at ambient blood glucose concentrations in the range 12.5±0.7 mmol/L for both VSG-treated and sham mice, which occurred at a single or multiple site across the islet but did not advance across the islet, were defined as “oscillations”. Increases that had a defined site of origin but did not spread across the full width of the imaged plane, were defined as “partial” waves (Fig. 2Bi, Supp. Mov. 1D). Those increases spreading across the whole islet were termed “waves” (Fig. 2Ai, 2Bii Supp. Mov. 1A, E). If the latter wave type was recurrent, we defined the behaviour as a “super wave” (Fig 2Biii, Supp. Mov. 1F). As illustrated in Fig. 2A, when imaged 0, 4 and 10 weeks after surgery, islets in sham-operated animals displayed a progressive loss of Ca^2+^ dynamics, as defined by the frequency and type of waves. Thus, when imaged at 0 weeks (Fig. 2Ai), wave behaviour (beginning at the bottom right; red area) area moved rapidly across the areas identified in yellow and blue. Comparable behaviour was seen at 4 weeks, with a similar site of origin of the wave (Fig. 2Aii) but was lost at 10 weeks post sham surgery, even though there was no significant weight difference between the two groups (Fig. 2Aiii, Supp. Mov. 1C).

**Figure 2:**
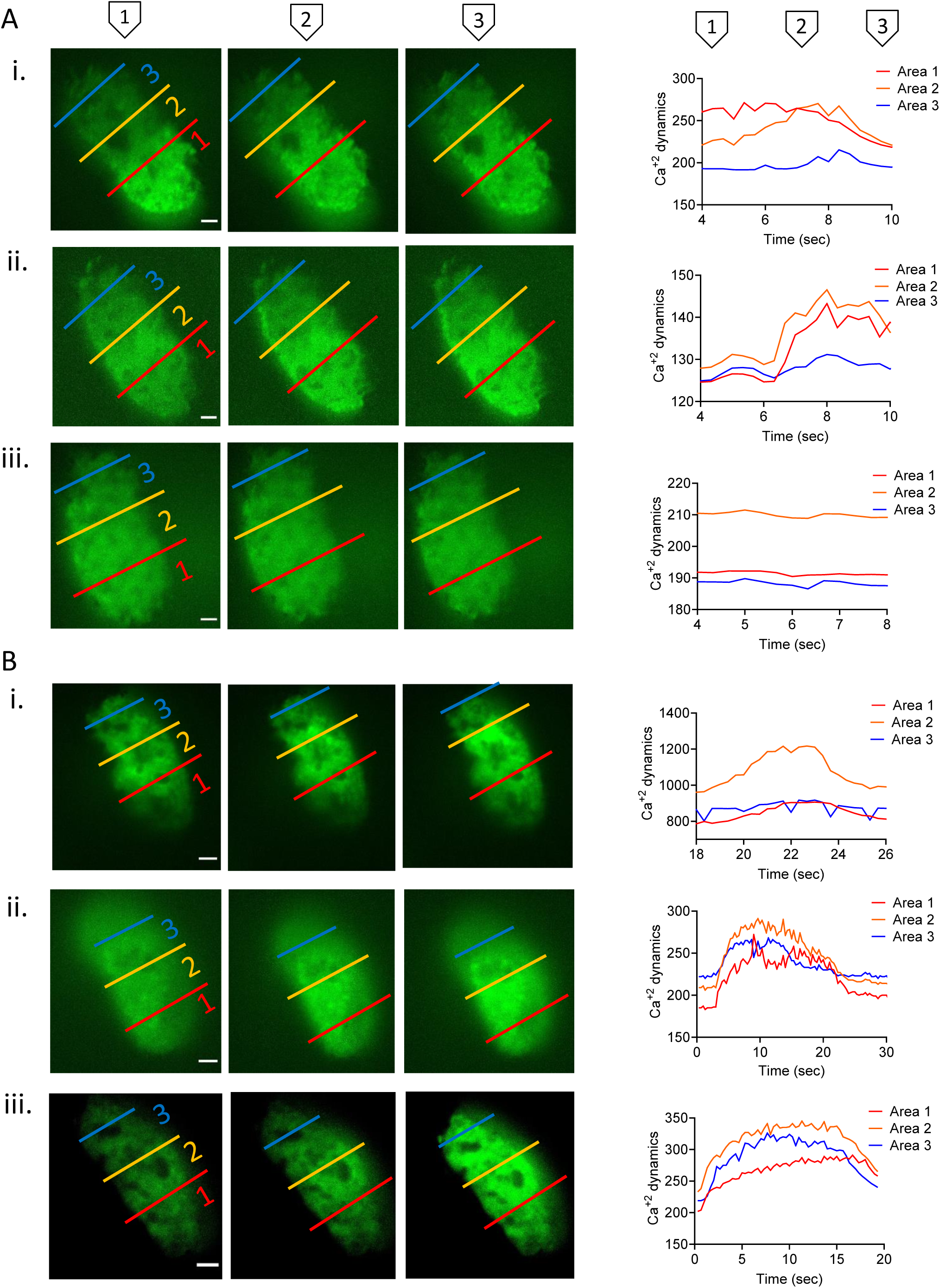
Description of Ins1Cre:GCaMPf^fl/fl^ islet Ca^+2^ dynamics: super wave, full wave, partial wave and no activity. Ins1Cre:GCaMPf^fl/fl^ islets implanted in the anterior chamber of the eye and imaged for 400 frames (133 secs) using a spinning disk confocal microscope (see Materials and Methods) at baseline week 0, postoperative week 4 and week 10. Each islet is separated in three Regions of Interest (ROIs) in order to categorise its activity. Red represents area 1 (distal islet), yellow represents area 2 (middle islet) and blue represents area 3 (proximal islet). Mean intensity is measured in each ROI for each frame (3 frames/ sec) and presented as Ca^+2^ dynamics. (A). Ins1Cre:GCaMPf^fl/fl^ islets implanted in a sham animal and imaged at (i) week 0 (full wave), (ii) 4 (full wave) and (iii) 10 (inactive). B (i) Ins1Cre:GCaMPf^fl/fl^ islet implanted in a VSG-treated animal (i) at week 0 (partial wave), (ii) week 4 (full wave) and (iii) week 10 (super wave). Scale: 100μm. Plasma glucose levels during imaging were 12.5±0.7 mmol/L.

In contrast, islets implanted into mice subject to VSG displayed sustained or gradually improving Ca^2+^ dynamics following surgery. Thus, the islet shown in Fig. 2Bi initially showed partial wave activity but progressed to full wave activity by week 4 (Fig. 2Bii, Supp. Mov. 1D) and to super wave by week 10 (Fig. 2Biii, Supp. Mov. 1F). A similar progression was seen for eight islets in three separate mice subjected to VSG (Fig. 3A, Supp. Fig. 3), whilst in six islets in three sham-operated mice a decline in behaviour was apparent after surgery (Fig. 3A). Remarkably, almost all islets transplanted into VSG animals displayed either wave or superwave behaviour by week eight, even if VSG-treated animals did not display further weight loss. This is significantly higher when compared to sham mice at the same timepoint (p=0.02) (Fig. 3A). By week 10, the activity of all sham-transplanted islets dropped to almost zero (p=0.004) (Fig. 3A). Mean wave front velocity, a measure of the speed of the wave calculated by distance (μm) divided by time (sec), across the islet was not different between groups at any time point explored. Similarly, no differences were apparent between wave velocities for the different wave types in either VSG or sham operated (Fig. 3B).

**Figure 3:**
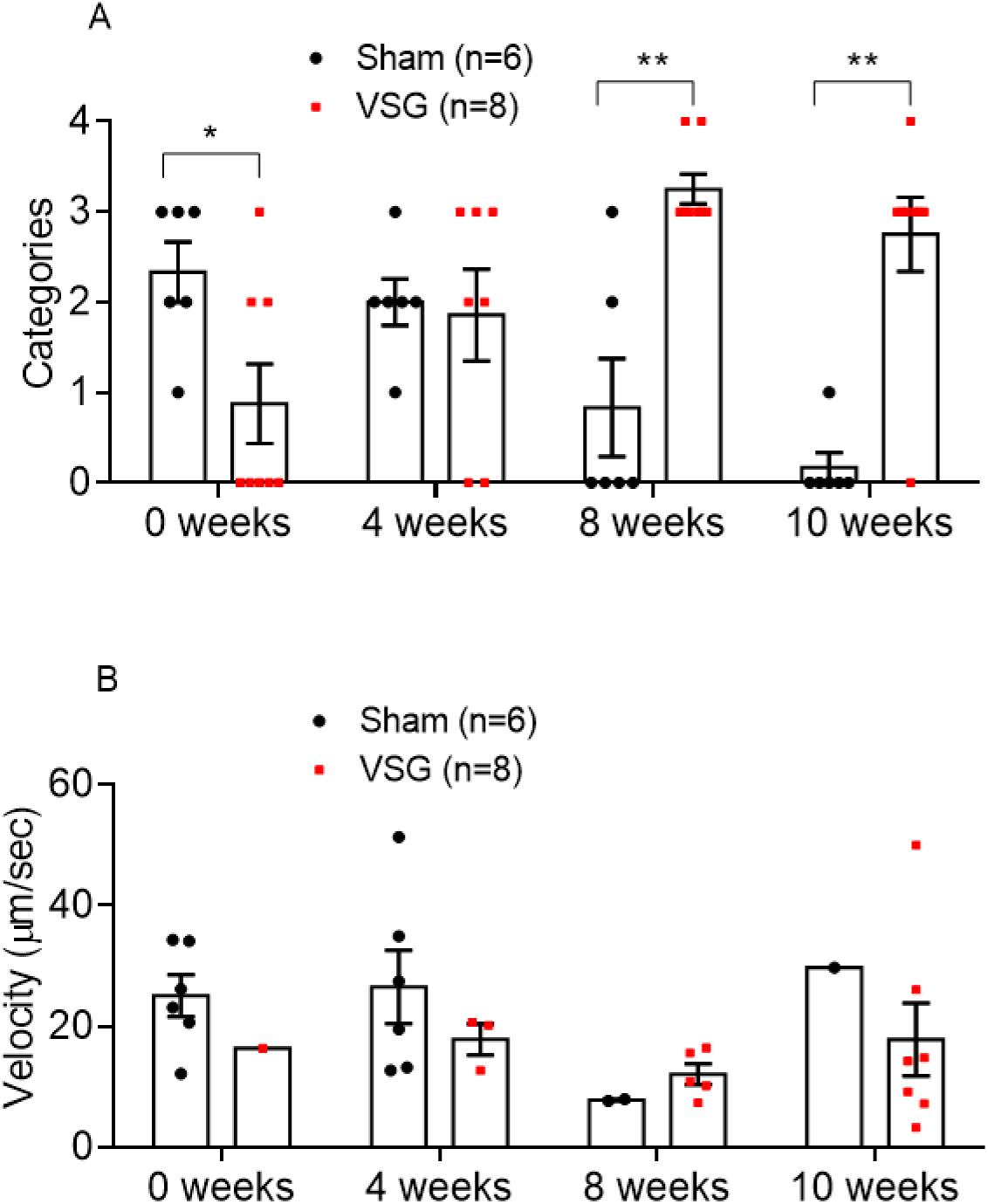
Ca^2+^ dynamics of Ins1Cre:GCaMPf^fl/fl^ islets in Sham and VSG-treated animals. (A). Categorisation of Ins1Cre:GCaMPf^fl/fl^ islets in sham (n=3 animals, n=6 islets) and VSG (n=3 animals, n=8 islets) mice on weeks 0 (baseline), 4, 8 and 12. Categories: 1. No activity, 2. Oscillations, 3. Partial Wave, 4. Wave, 5. Super Wave (B). Velocity of waves and partial waves of Ins1Cre:GCaMPf islets in sham and VSG animals calculated by d/ Δt and measured as μm/ sec. *P<0.05, **p<0.01, by Student t-test. Data are expressed as means ± SEM.

### Vertical Sleeve Gastrectomy maintains the number and strength of β cell – β cell connections

Coordinated activity of β cells is a feature of the healthy islet, and is likely to be important for the regulation of pulsatile insulin secretion (29). As shown in Fig. 4A and B, Pearson correlation analysis revealed no differences in apparent connectivity at week 0 (prior to surgery), whereas a progressive decline in connectivity was observed in the sham group. The number of connected cells (Fig. 4B), or the mean connectivity strength (R) (Fig. 4C, D) remained relatively constant in the VSG group, such that by week 10 these islets displayed significantly greater connectivity than the sham group (Fig. 4B). In summary, glucose-related Ca^2+^ signalling in VSG mice was characterized by higher magnitude and higher sensitivity to glucose when compared with sham mice, suggesting changes in glucose metabolism in the islets following VSG.

**Figure 4:**
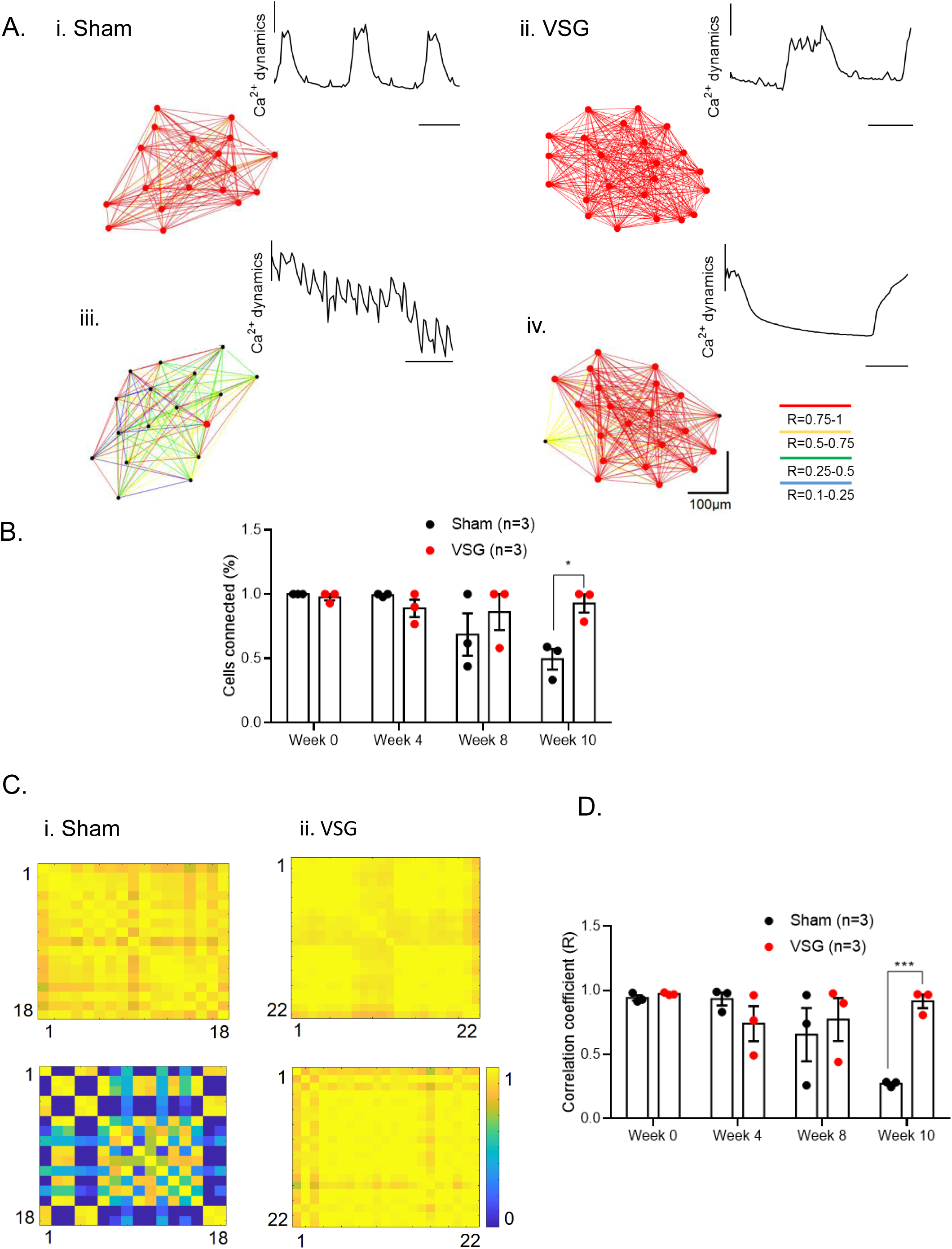
Cartesian functional connectivity and correlation coefficiency of islets before and after VSG or Sham surgery. (A) Cartesian functional connectivity maps displaying the correlation coefficients of β cells within the x-y position of analysed cells (dots). Cells are connected with a line where the strength of each cell pair correlation (the Pearson R statistic) is colour coded: red for R of 0.76 to 1.0, yellow for R of 0.51 to 0.75, green for R of 0.26-0.5 and blue for R of 0.1 to 0.25. Red dots represent β cells with the highest number of connected cell pairs. Ca^2+^ activity detected during the 30 sec (100 frames) imaging period analysed is displayed at the top right of each connectivity map. (B) The percentage of significantly connected cell pairs decreased significantly in the sham group at week 10. (C) Representative heatmaps depicting connectivity strength (Pearson R correlation) of all β cell pairs (x-y axis) presented in (A) (R values colour coded from 0 to 1, blue to yellow respectively). Yellow represents β cell pairs with high connectivity strength. (D) The average of correlation coefficient (R) decreased significantly in the sham group at week 10. N= 3 animals (1 islet/ animal). Data are means ± SEM and *p<0.05, **p<0.01, ***p<0.001 following 1-way ANOVA.

## Discussion

Using an intravital imaging approach developed in recent years to monitor islet function *in vivo* (26, 27), we provide here evidence that VSG causes a dramatic improvement in β cell Ca^2+^ dynamics, a useful assay of normal cellular function and proxy for insulin secretion (22, 30, 31). The use of such an approach addresses the challenges in dissecting the relative importance of the actions of bariatric surgery in changes observed in: (a) pancreatic insulin output versus peripheral insulin sensitivity, (b) β cell function versus mass, and (c) the time courses of changes, post-surgery.

Critically, we demonstrate that at similar, stimulatory glucose concentrations, islet Ca^2+^ dynamics and connectivity are dramatically increased in VSG versus sham-operated animals. Our data provide the first evidence we are aware of that alterations in β cell function occur both at the level of individual cells and across the islet ensemble after surgery, and are thus likely to play a pivotal role in improving insulin output. Changes in both β cell identity, reflecting altered gene expression (32-34), and in coordinated β cell activity across the islet, are important features of T2D (25, 35). The normalization of either thus presents an attractive therapeutic route towards improving insulin secretion in this disease. Importantly, whilst several studies have demonstrated changes in islet gene expression in rodent models related to hyperglycaemia and diabetes progression, such as obese diabetic (ZDF) rats and HFHSD mice (36, 37), few have examined the potential for reversing these changes as a therapeutic strategy (37, 38).

Central to the present study has been the use of VSG in obese mice as a model of human bariatric surgery (39, 40). Roux-en-Y Gastric Bypass (RYGB) and VSG are routinely deployed as an approach to treat human obesity, and both cause similar rates of T2D remission within the first post-operative year in man (41). In mice, VSG leads to initial rapid weight loss followed by a weight regain, unlike RYGB, but sustains improved glucose tolerance while offering a more tractable approach, with lower mortality (42, 43). Importantly, our study had a ten-week post-operative follow up and, by week eight, there was no significant weight difference between sham and VSG group. This allowed us to separate marked improvements in insulin secretion from significant weight loss without the need to pair-feed the sham group (43). Moreover, it corresponds with our previous findings in lean VSG-treated mice that demonstrated no weight difference when compared to sham mice at four weeks post-operatively, yet displayed improved glucose tolerance and corresponding insulin secretion curves during an IPGTT (44). Insulin tolerance tests at week eight demonstrated that VSG mice had improved insulin sensitivity, an effect previously attributed in humans to rapid and significant enhancement of post-operative hepatic insulin clearance (45). It has recently been suggested that, in animal models of surgery, hepatic insulin clearance is related to peripheral, rather than hepatic, insulin sensitivity (46).

Accelerated glucose clearance in VSG-treated animals was accompanied by increased insulin secretion in response to glucose at 15 and 30 min., consistent with previous studies using VSG models (39, 47, 48). The fact that the insulin response to IPGTT was equally robust suggests that this effect is not solely due to an increased spike in blood glucose associated with elevated gastric emptying rates and upregulated glucose absorption, as has been previously postulated (9, 49). Increased insulin secretion in the face of lower plasma glucose demonstrates enhanced β cell glucose sensitivity, consistent with cell-autonomous changes in islet function, alterations in circulating levels of other regulators of secretion, or an increase in β cell number. Analysis of the endogenous pancreatic β cell mass showed no increase in VSG versus sham-operated mice, pointing to a functional change rather than a change in endogenous β cell mass, as underlying increased insulin output. Moreover, and though this could not be quantitated accurately due to the lack of focal distances stacking data, we saw no evidence for a change in the β cell mass of islets engrafted into the eye. This is in line with previous findings demonstrating that there is no islet hyperplasia or increased β cell turnover following bariatric surgery in humans or rats with obesity (50, 51). These findings contrast other studies that have reported -over similar times scales-increasing (52-54) or decreasing (48, 55) β cell mass differences, which may reflect pre-operative metabolic state or other factors.

Given many reports of increased GLP-1 release after bariatric surgery in both humans (56) and rodents (42, 57), here we explored levels of both circulating fasting and post-glucose gavage GLP-1 levels. Importantly, the peak in GLP-1 following oral gavage did not differ between sham and VSG mice, indicating that an enhanced insulinotropic effect of the incretin is unlikely to explain dramatic increase in insulin secretion observed. Furthermore, enhanced insulin secretion was seen in mice treated with VSG even during IPGTT, where the stimulation of GLP-1 secretion is negligible.

Interestingly, VSG-treated mice did display significantly higher circulating GLP-1 levels under basal (fasting) conditions. Apart from increasing glucose-stimulated insulin secretion and enhancing insulin gene transcription (58, 59), GLP-1 also inhibits β cell apoptosis in animal models of diabetes (60, 61). Thus, although the underlying mechanisms remain unclear, increased basal GLP-1 levels might provide a partial explanation for the enhanced responses to glucose and the euglycemic effects observed after surgery. A number of studies have shown that, in post-operative patients with T2D, GLP-1 receptor (GLP-1R) blockade with the GLP-1R antagonist Exendin-(9-39) causes significant reduction of insulin secretion when compared to a control group with lower GLP-1 levels, indicating an effect on β cell function (62, 63). Nonetheless, other studies have demonstrated that blocking GLP-1R in bariatric patients impairs glucose tolerance but not to a greater degree than before surgery, or when compared to non-operated patients (64, 65). Furthermore, Ye and colleagues found that pharmacological or genetic blockade or elimination of GLP-1R signaling in rats or mice, respectively, had no impact on the ability of RYGB to lower body weight (66, 67). Taken together, these earlier data suggest that GLP-1 signaling may not be the main mediator of T2D remission but is likely to contribute (68). Changes in the levels of other circulating factors are thus likely to be involved in the apparent increase in β cell function. Lowered levels of circulating lipids (69), inflammatory cytokines (70), bile acids (71), glucocorticoids (72) or microbiome-derived products (73) are potential candidates.

An important aspect of the present study has been to examine, at the cellular level, the functional connectivity between β cells before and after VSG or sham surgery. The percentage of significantly-connected cell pairs and correlation coefficient decreased substantially in the sham group at week ten, while in the VSG group these parameters remained stable for the duration of the study. We would note that hub/follower behaviour (i.e. the existence of a “power law” in the degree of connectedness) (23) was not readily apparent in the present study. More rapid acquisition rates are likely to be needed to reveal such a hierarchy. Furthermore, we note that wave-like behaviour is more often apparent in the islet *in vivo* using GCaMP6f as the Ca^2+^ sensor (26) than in some of our own and others’ earlier studies (23, 25) using entrapped, synthetic Ca^2+^ probes. Nonetheless, clusters of apparent “leader” β cells, corresponding to the point at which a rise in Ca^2+^ was first observed at the beginning of a wave, were easily identified in many cases (e.g. Fig. 2A), and these have previously been reported (23) to correspond to the hub cell population. Interestingly, the origin of the waves was similar within a given islet assayed several weeks apart, indicating that the cells which initiate them (leaders) represent a stable population, at least over the time frame (≤ 10 weeks) studied here.

Taken together, our results indicate that bariatric surgery improves glycaemic control at least partially by maintaining: (a) functional β cell identity and (b) coordinated activity across the islet. A possible explanation for our data may be that the improvement in islet function follows the improvement in glycaemia. However, since insulin sensitivity was barely altered by VSG, it is unclear whether extrapancreatic events could be the drivers for improved islet function and insulin output.

Although the surgical model used provides us with novel information on β cell activity via continuous monitoring, there are undoubtedly limitations in the use of islets engrafted into the ACE. These include potential differences between the vascularization and innervation at this site compared to pancreatic in situ islets (74). Our findings on ACE-engrafted islet reactivation following VSG are however in line with previous results focusing on pancreatic islets isolated postmortem. Thus, Douros et al (43) performed Ca^2+^ imaging *in vitro* in mouse islets postmortem and showed that the percentage of islets displaying Ca^2+^ oscillations in response to glucose was enhanced 2.2-fold in the VSG group, indicating increased islet glucose sensitivity after surgery. In addition, VSG altered the islet transcriptome, affecting genes involved in insulin secretion and Ca^2+^ signaling (43). However, these earlier studies were cross-sectional in nature, and as such did not explore the apparent reactivation *in vivo* of individual islets and β cells in the living animal, as described here.

In conclusion, our findings provide further evidence for the protective effect of bariatric surgery in T2D, irrespective of weight loss, and demonstrate direct effects on β-cell function and coordination in the living animal. Future challenges are to understand more fully the mechanisms through which these changes are affected at the paracrine, endocrine and cellular levels.

## Methods

### Animals

All animal procedures undertaken were approved by the British Home Office under the UK Animal (Scientific Procedures) Act 1986 (Project License PPL PA03F7F07 to I.L.) with approval from the local ethical committee (Animal Welfare and Ethics Review Board, AWERB), at the Central Biological Services (CBS) unit at the Hammersmith Campus of Imperial College London.

Adult male C57BL/6J mice (Envigo, Huntingdon U.K.) were maintained under controlled temperature (21-23°C) and light (12:12 hr light-dark schedule, lights on at 0700). From the age of 8 weeks they were put on a 58 kcal% Fat and Sucrose diet (D12331, Research Diet, New Brunswick, NJ) ad libitum to induce obesity and diabetes. Four weeks after the start of this diet, the animals underwent islet transplantation into the anterior chamber of the eye of genetically modified islets expressing GCaMP6f to allow for intravital measurements of cytosolic Ca^2+^. Four weeks after islet transplantation, the animals underwent either a vertical sleeve gastrectomy or a sham surgery as described below.

Ins1Cre:GCaMPf^fl/fl^ mice, *used as donors for islet transplantation* were generated by crossing crossed Ins1Cre mice (provided by J Ferrer, this Department) to mice that express GCaMP6f downstream of a LoxP-flanked STOP cassette (The Jackson Laboratory, stock no. 028865). Islets donated from either sex were used for transplantation.

### Islet transplantation into the anterior chamber of the mouse eye (ACE)

Pancreatic islets were isolated and cultured as described previously (75). For transplantation, 10-20 islets were aspirated with a 27-gauge blunt eye cannula (BeaverVisitec, UK) connected to a 100ul Hamilton syringe (Hamilton) via 0.4-mm polyethylene tubing (Portex Limited). Prior to surgery, mice were anaesthetised with 2-4% isoflurane (Zoetis) and placed in a stereotactic frame to stabilise the head. The cornea was incised near the junction with the sclera, being careful not to damage the iris. Then, the blunt cannula, pre-loaded with islets, was inserted into the ACE and islets were expelled (average injection volume 20 µl for 10 islets). Carprofen (Bayer, UK) and eye ointment were administered post-surgery.

### Vertical Sleeve Gastrectomy

Three days before bariatric or sham surgery, animals were exposed to liquid diet (20% dextrose) and remained on this diet for up to four days post operatively. Following this, mice were returned to high fat/high sucrose diet until euthanasia and tissues harvested ten weeks post bariatric surgery. Anaesthesia was induced and maintained with isoflurane (1.5-2%). A laparotomy incision was made, and the stomach was isolated outside the abdominal cavity. A simple continuous pattern of suture extending through the gastric wall and along both gastric walls was placed to ensure the main blood vessels were contained. Approximately 60% of the stomach was removed, leaving a tubular remnant. The edges of the stomach were inverted and closed by placing two serosae only sutures, using Lembert pattern. The initial full thickness suture was subsequently removed. Sham surgeries were performed by isolating the stomach and performing a 1 mm gastrotomy on the gastric wall of the fundus. All animals received a five-day course of SC antibiotic injections (Ciprofloxacin 0.1mg/kg).

### In vivo Ca^2+^ imaging of Ins1Cre:GCaMPf^fl/fl^ islets in the ACE

A minimum of four weeks was allowed for full implantation of islets before imaging. Imaging sessions were performed as previously described (26) with the mouse held in a stereotactic frame and the eye gently retracted, with the animal maintained under 2-4% isoflurane anaesthesia. All imaging experiments were conducted using a spinning disk confocal microscope (Nikon Eclipse Ti, Crest spinning disk, 20x water dipping 1.0 NA objective). The signal from GCaMP6f fluorophore (ex. 488 nm, em. 525±25 nm) was monitored in time-series experiments for up to 20 min. at a rate of 3 frames/ sec. Ca^2+^ traces were recorded for three min, with a mean blood glucose reading (across six islets in three separate animals per group) of 12.5±0.7. mmol/L. Islets were continuously monitored, and the focus was manually adjusted to counteract movement. Animals were imaged 3 days prior to Vertical Sleeve Gastrectomy (baseline) and then at four, eight and ten weeks post-operatively.

### Glucose Tolerance Tests

Mice were fasted overnight (total 16 h) and given free access to water. At 0900, glucose (3 g/kg body weight) was administered via intraperitoneal injection or oral gavage. Blood was sampled from the tail vein at 0, 5, 15, 30, 60 and 90 min. after glucose administration. Blood glucose was measured with an automatic glucometer (Accuchek; Roche, Burgess Hill, UK).

### Insulin Tolerance Tests

Mice were fasted for 8 h and given free access to water. At 1500, human insulin (Actrapid, Novo Nordisk) (1.5U/kg body weight) was administered via intraperitoneal injection. Blood was sampled from the tail vein at 0, 15, 30, 60 and 90 min after insulin administration. Blood glucose was measured with an automatic glucometer (Accuchek; Roche, Burgess Hill, UK).

### Plasma insulin and GLP-1 measurement

To quantify circulating insulin and GLP-1(1-37) levels, 100μl of blood was collected from the tail vein into heparin-coated tubes (Sarstedt, Beaumont Leys, UK). Plasma was separated by sedimentation at 10,000***g*** for 10 min. (4°C). Plasma insulin levels were measured in 5μl aliquots and GLP-1(1-37) levels were measured in 10μl aliquots by ELISA kits from Crystal Chem (USA).

### Immunohistochemistry of pancreas sections

Isolated pancreata were fixed in 10% (vol/vol) buffered formalin and embedded in paraffin wax within 24 h of removal. Slides (5 μm) were submerged sequentially in Histoclear (Sigma, UK) followed by washing in decreasing concentrations of ethanol to remove paraffin wax. Permeabilised pancreatic slices were blotted with ready-diluted anti-guinea pig insulin (Agilent Technologies, USA) and anti-mouse glucagon (Sigma, UK) primary antibody (1:1000). Slides were visualised by subsequent incubation with Alexa Fluor 488 and 568-labelled donkey anti-guinea pig and anti-mouse antibody. Samples were mounted on glass slides using VectashieldTM (Vector Laboratories, USA) containing DAPI. Images were captured on a Zeiss Axio Observer.Z1 motorised inverted widefield microscope fitted with a Hamamatsu Flash 4.0 Camera using a Plan-Apochromat 206/0.8 M27 air objective with Colibri.2 LED illumination. Data acquisition was controlled with Zeiss Zen Blue 2012 Software. Fluorescence quantification was achieved using Image J (https://imagej.nih.gov/ij/). Whole pancreas was used to quantitate cell mass.

### Statistical Analysis

Data were analysed using GraphPad PRISM 7.0 software. Significance was tested using unpaired Student’s two-tailed t-tests with Bonferroni post-tests for multiple comparisons, or two-way ANOVA as indicated. P<0.05 was considered significant and errors signify ± SEM.

### Pearson (R)-based connectivity and correlation analyses

Correlation analyses between the Ca^2+^ signal time series for all cell pairs in an imaged islet were performed in MATLAB using a modified custom-made script (26). B cell intensity was measured in 100 consecutive frames (30 sec.). Regions of Interest (ROI, 18-40 per islet, depending on size) were selected with single or near single cell resolution (i.e. 10-20μm diameter) and captured approximately 95% of the fluorescence of the image plane. Data were smoothed using a retrospective averaging method (previous 10 values). The correlation function R between all possible (smoothed) cell pair combinations (excluding the autocorrelation) was assessed using Pearson’s correlation. The Cartesian co-ordinates of the imaged cells were then incorporated in the construction of connectivity line maps. Cell pairs were connected with a straight line, the colour of which represented the correlation strength and was assigned to a colour-coded light-dark ramp (R=0.1-0.25 [blue], 0.26-0.5 [green], R=0.51-0.75 [yellow], R=0.76-1.0 [red]). Cells with the highest number of possible cell pair combinations are shown in red. Data are also displayed as heatmap matrices, indicating individual cell pair connections on each axis (min. = 0; max. = 1). The positive R values (excluding the auto-correlated cells) and the percentage of cells that were significantly connected to one another were averaged and compared between groups.

## Funding

E.A. was supported by a grant from the Rosetrees Trust (M825) and from the British Society for Neuroendocrinology. G.R., E.G. and P.C. were supported by a Wellcome Trust Investigator (212625/Z/18/Z) Award, MRC Programme grants (MR/R022259/1, MR/J0003042/1, MR/L020149/1), and Experimental Challenge Grant (DIVA, MR/L02036X/1), MRC (MR/N00275X/1), and Diabetes UK (BDA/11/0004210, BDA/15/0005275, BDA 16/0005485) grants. This project has received funding from the European Union’s Horizon 2020 research and innovation programme via the Innovative Medicines Initiative 2 Joint Undertaking under grant agreement No. 115881 (RHAPSODY) to G.R. V.S. and K.S. were supported by Harry Keen Diabetes UK Fellowship (BDA 15/0005317). I.L. is supported by a project grant from Diabetes UK (16/0005485).

## Supplementary data

**Figure 1:**
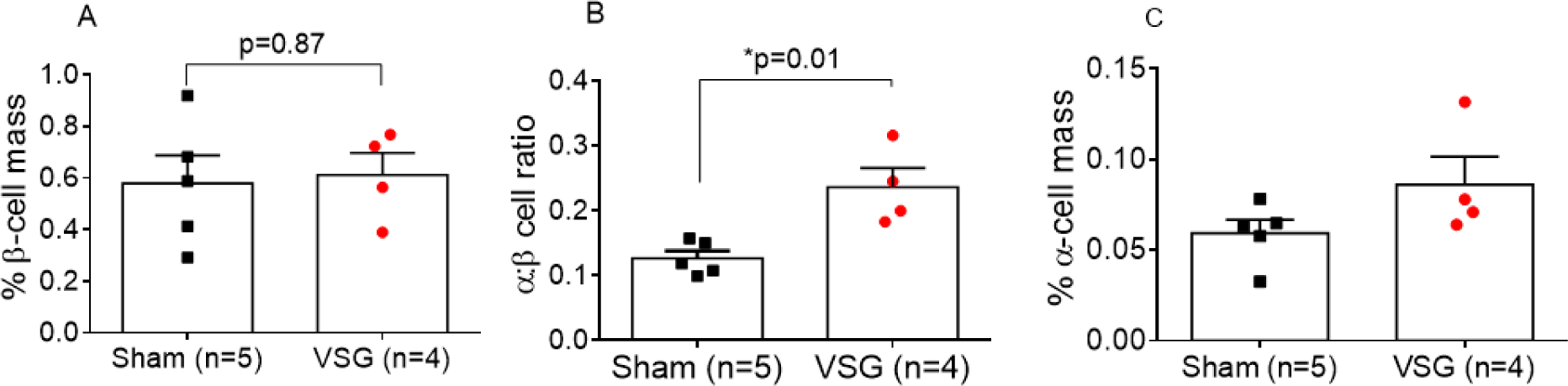
Percentage of pancreatic area occupied by and ratio of α to β cells. (A) Percentage of pancreatic area occupied by β cells measured by immunofluorescent staining of pancreatic islets using anti-insulin antibody against total pancreatic area. (B) Ratio of α to β cells in pancreatic islets of VSG (n=4) and Sham (n=5) mice measured by immunofluorescent staining of pancreatic islets using anti-glucagon (red) and anti-insulin (green) antibody. (C) Percentage of pancreatic area occupied by α cells measured by immunofluorescent staining of pancreatic islets using anti-glucagon antibody against total pancreatic area. Each point represents an average of all islets (10-30/ slide) present in three permeabilised pancreatic slices, separated by 600μm. Total pancreatic area is measured in each slide and percentage is calculated accordingly.*P<0.05, by unpaired Student’s t-test. Data are expressed as means ± SEM.

**Figure 2:**
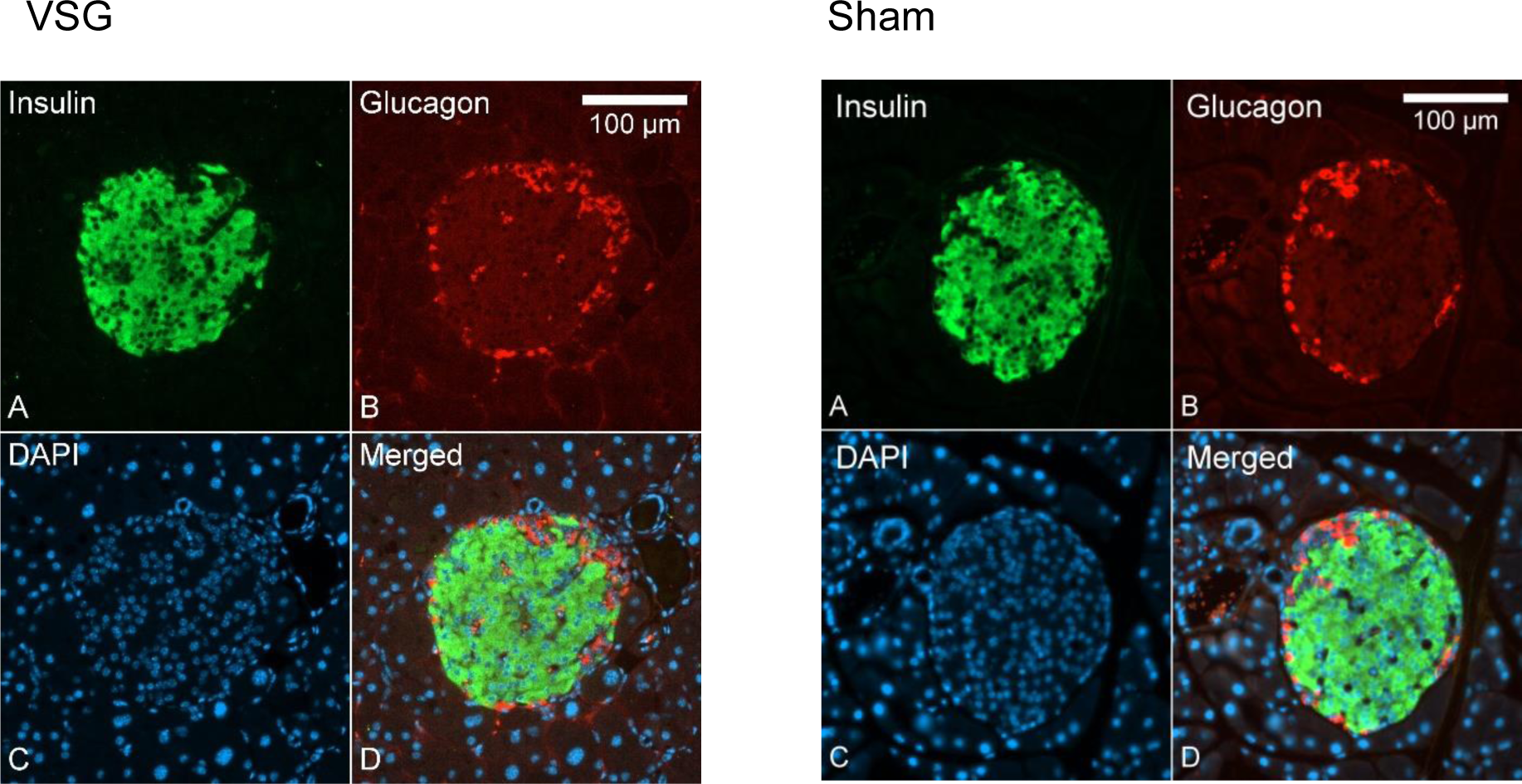
Immunofluorescence staining of pancreatic islets. Anti-glucagon (red) and anti-insulin (green) antibodies where used in VSG (n=4) and Sham (n=5) mice.

**Figure 3:**
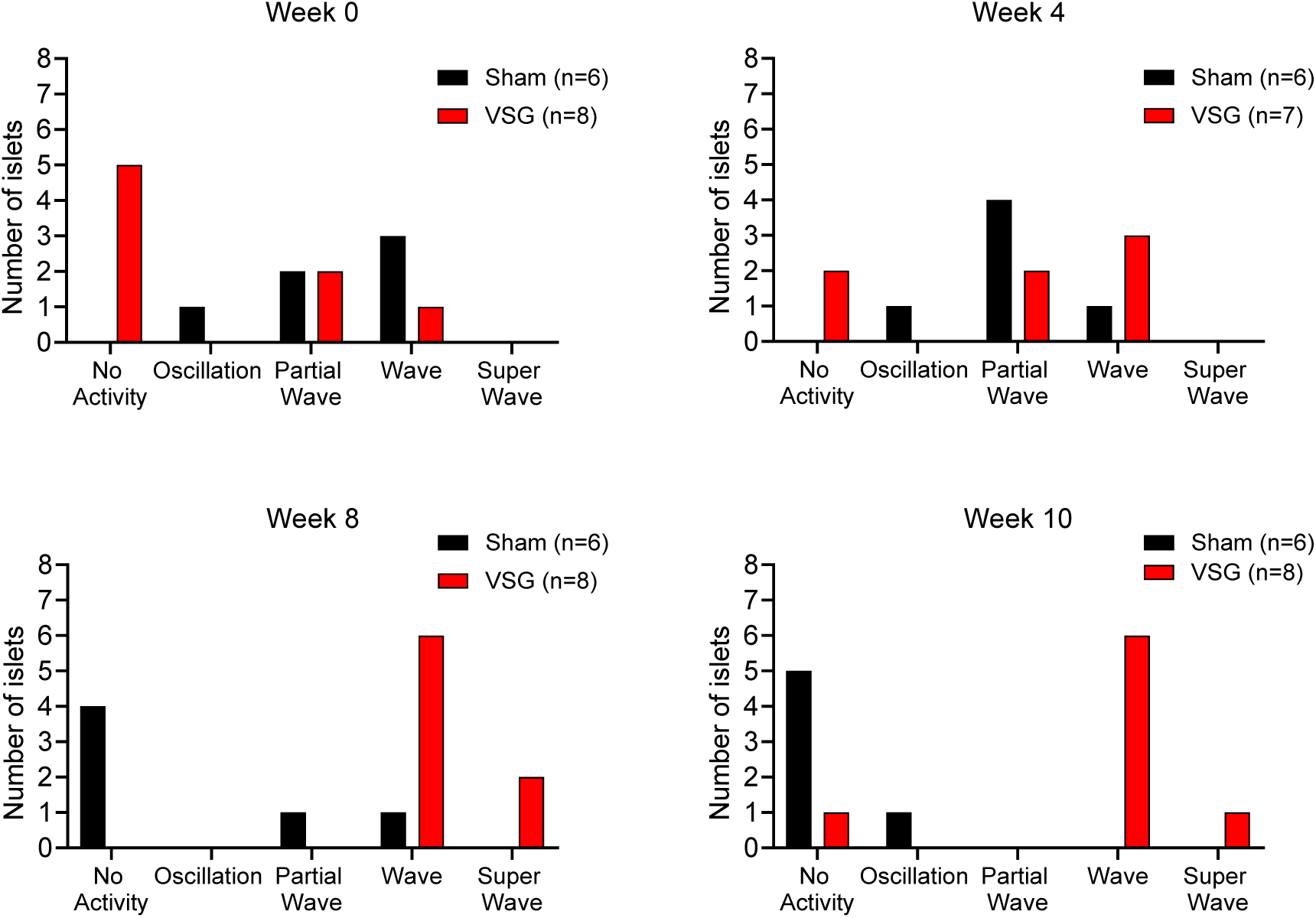
Ca^2+^ dynamics of Ins1Cre:GCaMPf^fl/fl^ islets in Sham and VSG-treated animals. Grouped categorisation of Ins1Cre:GCaMPf islets in sham (n=3 animals, n=6 islets) and VSG (n=3 animals, n=8 islets) mice on weeks 0 (baseline), 4, 8 and 12.

Movie 1:

**Description of Ins1Cre:GCaMPf^fl/fl^ islet when in full wave, partial wave and no activity.** (A) Ins1Cre:GCaMPf^fl/fl^ islet implanted in a sham animal, imaged using a spinning disk confocal microscope at week 0 (full wave), (B) 4 (full wave) and (C) 12 (inactive). (D) Ins1Cre:GCaMPf^fl/fl^ islet implanted in VSG-treated animal, imaged at week 0 (partial wave), (E) 4 (full wave) and (F) 12 (super wave). Plasma glucose levels during imaging were 12.5±0.7 mmol/L measured at 5 min intervals.

## Notes

### Competing Interest Statement

GA Rutter has received grant funds from Servier Laboratories and Sun Pharmaceutical Industries Ltd. These funders were not involved in any of the studies discussed here. The remaining authors have no conflict of interest to disclose.

## References

1. Anonymous (2017) National Institute of Diabetes and Digestive and Kidney Diseases: National Diabetes Statistics Report.

2. Anonymous (2019) Diabetes UK: Diabetes Facts and stats

3. K. D. Hall, S. Kahan, Maintenance of Lost Weight and Long-Term Management of Obesity. Med Clin North Am 102, 183–197 (2018).

4. M. J. Franz, J. L. Boucher, S. Rutten-Ramos, J. J. VanWormer, Lifestyle weight-loss intervention outcomes in overweight and obese adults with type 2 diabetes: a systematic review and meta-analysis of randomized clinical trials. J Acad Nutr Diet 115, 1447–1463 (2015).

5. K. Abegg et al., Effect of bariatric surgery combined with medical therapy versus intensive medical therapy or calorie restriction and weight loss on glycemic control in Zucker diabetic fatty rats. Am J Physiol Regul Integr Comp Physiol 308, R321–329 (2015).

6. A. A. Gumbs, I. M. Modlin, G. H. Ballantyne, Changes in insulin resistance following bariatric surgery: role of caloric restriction and weight loss. Obes Surg 15, 462–473 (2005).

7. S. R. Kashyap, D. L. Bhatt, P. R. Schauer, S. Investigators, Bariatric surgery vs. advanced practice medical management in the treatment of type 2 diabetes mellitus: rationale and design of the Surgical Therapy And Medications Potentially Eradicate Diabetes Efficiently trial (STAMPEDE). Diabetes Obes Metab 12, 452–454 (2010).

8. N. Q. Nguyen et al., Upregulation of intestinal glucose transporters after Roux-en-Y gastric bypass to prevent carbohydrate malabsorption. Obesity (Silver Spring) 22, 2164–2171 (2014).

9. G. Baud et al., Sodium glucose transport modulation in type 2 diabetes and gastric bypass surgery. Surg Obes Relat Dis 12, 1206–1212 (2016).

10. P. Dadson et al., Brown adipose tissue lipid metabolism in morbid obesity: Effect of bariatric surgery-induced weight loss. Diabetes Obes Metab 20, 1280–1288 (2018).

11. P. Dadson et al., Effect of Bariatric Surgery on Adipose Tissue Glucose Metabolism in Different Depots in Patients With or Without Type 2 Diabetes. Diabetes Care 39, 292–299 (2016).

12. H. Immonen et al., Effect of bariatric surgery on liver glucose metabolism in morbidly obese diabetic and non-diabetic patients. J Hepatol 60, 377–383 (2014).

13. C. Dirksen et al., Exaggerated release and preserved insulinotropic action of glucagon-like peptide-1 underlie insulin hypersecretion in glucose-tolerant individuals after Roux-en-Y gastric bypass. Diabetologia 56, 2679–2687 (2013).

14. A. P. Chambers et al., Similar effects of roux-en-Y gastric bypass and vertical sleeve gastrectomy on glucose regulation in rats. Physiol Behav 105, 120–123 (2011).

15. S. Al-Sabah et al., Incretin response to a standard test meal in a rat model of sleeve gastrectomy with diet-induced obesity. Obes Surg 24, 95–101 (2014).

16. M. Nannipieri et al., The role of beta-cell function and insulin sensitivity in the remission of type 2 diabetes after gastric bypass surgery. J Clin Endocrinol Metab 96, E1372–1379 (2011).

17. G. M. Campos et al., Improvement in peripheral glucose uptake after gastric bypass surgery is observed only after substantial weight loss has occurred and correlates with the magnitude of weight lost. J Gastrointest Surg 14, 15–23 (2010).

18. K. N. Bojsen-Moller et al., Early enhancements of hepatic and later of peripheral insulin sensitivity combined with increased postprandial insulin secretion contribute to improved glycemic control after Roux-en-Y gastric bypass. Diabetes 63, 1725–1737 (2014).

19. A. Mallipedhi et al., Temporal changes in glucose homeostasis and incretin hormone response at 1 and 6 months after laparoscopic sleeve gastrectomy. Surg Obes Relat Dis 10, 860–869 (2014).

20. S. A. Brethauer et al., Early effects of gastric bypass on endothelial function, inflammation, and cardiovascular risk in obese patients. Surg Endosc 25, 2650–2659 (2011).

21. K. Samaras, A. Viardot, N. K. Botelho, A. Jenkins, R. V. Lord, Immune cell-mediated inflammation and the early improvements in glucose metabolism after gastric banding surgery. Diabetologia 56, 2564–2572 (2013).

22. D. Laurent et al., Pancreatic beta-cell imaging in humans: fiction or option? Diabetes Obes Metab 18, 6–15 (2016).

23. N. R. Johnston et al., Beta Cell Hubs Dictate Pancreatic Islet Responses to Glucose. Cell Metab 24, 389–401 (2016).

24. R. K. Benninger, D. W. Piston, Cellular communication and heterogeneity in pancreatic islet insulin secretion dynamics. Trends Endocrinol Metab 25, 399–406 (2014).

25. D. J. Hodson et al., Lipotoxicity disrupts incretin-regulated human beta cell connectivity. J Clin Invest 123, 4182–4194 (2013).

26. V. Salem et al., Leader beta-cells coordinate Ca2+ dynamics across pancreatic islets in vivo. Nature Metabolism 1, 615–629 (2019).

27. S. Speier et al., Noninvasive in vivo imaging of pancreatic islet cell biology. Nat Med 14, 574–578 (2008).

28. M. S. Winzell, B. Ahren, The high-fat diet-fed mouse: a model for studying mechanisms and treatment of impaired glucose tolerance and type 2 diabetes. Diabetes 53 Suppl 3, S215–219 (2004).

29. G. A. Rutter, D. J. Hodson, Beta cell connectivity in pancreatic islets: a type 2 diabetes target? Cell Mol Life Sci 72, 453–467 (2015).

30. P. Gilon, H. Y. Chae, G. A. Rutter, M. A. Ravier, Calcium signaling in pancreatic beta-cells in health and in Type 2 diabetes. Cell Calcium 56, 340–361 (2014).

31. L. S. Satin, P. C. Butler, J. Ha, A. S. Sherman, Pulsatile insulin secretion, impaired glucose tolerance and type 2 diabetes. Mol Aspects Med 42, 61–77 (2015).

32. A. Segerstolpe et al., Single-Cell Transcriptome Profiling of Human Pancreatic Islets in Health and Type 2 Diabetes. Cell Metab 24, 593–607 (2016).

33. M. Solimena et al., Systems biology of the IMIDIA biobank from organ donors and pancreatectomised patients defines a novel transcriptomic signature of islets from individuals with type 2 diabetes. Diabetologia 61, 641–657 (2018).

34. G. D. Gutierrez, J. Gromada, L. Sussel, Heterogeneity of the Pancreatic Beta Cell. Front Genet 8, 22 (2017).

35. G. A. Rutter, T. J. Pullen, D. J. Hodson, A. Martinez-Sanchez, Pancreatic beta-cell identity, glucose sensing and the control of insulin secretion. Biochem J 466, 203–218 (2015).

36. L. E. Parton et al., Limited role for SREBP-1c in defective glucose-induced insulin secretion from Zucker diabetic fatty rat islets: a functional and gene profiling analysis. Am J Physiol Endocrinol Metab 291, E982–994 (2006).

37. R. Roat et al., Alterations of pancreatic islet structure, metabolism and gene expression in diet-induced obese C57BL/6J mice. PLoS One 9, e86815 (2014).

38. C. Dai et al., Stress-impaired transcription factor expression and insulin secretion in transplanted human islets. J Clin Invest 126, 1857–1870 (2016).

39. D. Garibay, B. P. Cummings, A Murine Model of Vertical Sleeve Gastrectomy. J Vis Exp 10.3791/56534 (2017).

40. J. D. Douros et al., Enhanced Glucose Control Following Vertical Sleeve Gastrectomy Does Not Require a beta-Cell Glucagon-Like Peptide 1 Receptor. Diabetes 67, 1504–1511 (2018).

41. R. Murphy et al., Laparoscopic Sleeve Gastrectomy Versus Banded Roux-en-Y Gastric Bypass for Diabetes and Obesity: a Prospective Randomised Double-Blind Trial. Obes Surg 28, 293–302 (2018).

42. P. Larraufie et al., Important Role of the GLP-1 Axis for Glucose Homeostasis after Bariatric Surgery. Cell Rep 26, 1399–1408 e1396 (2019).

43. J. D. Douros et al., Sleeve gastrectomy rapidly enhances islet function independently of body weight. JCI Insight 4 (2019).

44. E. Akalestou, L. Lopez-Noriega, I. Leclerc, G. A. Rutter, Vertical sleeve gastrectomy lowers kidney SGLT2 expression in the mouse. bioRxiv https://doi.org/10.1101/741330 (2019).

45. I. Malandrucco et al., Very-low-calorie diet: a quick therapeutic tool to improve beta cell function in morbidly obese patients with type 2 diabetes. Am J Clin Nutr 95, 609–613 (2012).

46. I. Asare-Bediako et al., Variability of Directly Measured First-Pass Hepatic Insulin Extraction and Its Association With Insulin Sensitivity and Plasma Insulin. Diabetes 67, 1495–1503 (2018).

47. D. M. Arble, D. A. Sandoval, F. W. Turek, S. C. Woods, R. J. Seeley, Metabolic effects of bariatric surgery in mouse models of circadian disruption. Int J Obes (Lond) 39, 1310–1318 (2015).

48. A. K. McGavigan et al., TGR5 contributes to glucoregulatory improvements after vertical sleeve gastrectomy in mice. Gut 66, 226–234 (2017).

49. J. B. Cavin et al., Differences in Alimentary Glucose Absorption and Intestinal Disposal of Blood Glucose After Roux-en-Y Gastric Bypass vs Sleeve Gastrectomy. Gastroenterology 150, 454–464 e459 (2016).

50. J. J. Meier, A. E. Butler, R. Galasso, P. C. Butler, Hyperinsulinemic hypoglycemia after gastric bypass surgery is not accompanied by islet hyperplasia or increased beta-cell turnover. Diabetes Care 29, 1554–1559 (2006).

51. W. B. Inabnet et al., The utility of [(11)C] dihydrotetrabenazine positron emission tomography scanning in assessing beta-cell performance after sleeve gastrectomy and duodenal-jejunal bypass. Surgery 147, 303–309 (2010).

52. M. E. Patti et al., Heterogeneity of proliferative markers in pancreatic beta-cells of patients with severe hypoglycemia following Roux-en-Y gastric bypass. Acta Diabetol 54, 737–747 (2017).

53. G. J. Service et al., Hyperinsulinemic hypoglycemia with nesidioblastosis after gastric-bypass surgery. N Engl J Med 353, 249–254 (2005).

54. S. Zhang et al., Increased beta-Cell Mass in Obese Rats after Gastric Bypass: A Potential Mechanism for Improving Glycemic Control. Med Sci Monit 23, 2151–2158 (2017).

55. B. P. Cummings et al., Bile-acid-mediated decrease in endoplasmic reticulum stress: a potential contributor to the metabolic benefits of ileal interposition surgery in UCD-T2DM rats. Dis Model Mech 6, 443–456 (2013).

56. C. R. Hutch, D. Sandoval, The Role of GLP-1 in the Metabolic Success of Bariatric Surgery. Endocrinology 158, 4139–4151 (2017).

57. D. Garibay et al., beta Cell GLP-1R Signaling Alters alpha Cell Proglucagon Processing after Vertical Sleeve Gastrectomy in Mice. Cell Rep 23, 967–973 (2018).

58. P. E. MacDonald et al., The multiple actions of GLP-1 on the process of glucose-stimulated insulin secretion. Diabetes 51 Suppl 3, S434–442 (2002).

59. G. Skoglund, M. A. Hussain, G. G. Holz, Glucagon-like peptide 1 stimulates insulin gene promoter activity by protein kinase A-independent activation of the rat insulin I gene cAMP response element. Diabetes 49, 1156–1164 (2000).

60. M. Cornu et al., Glucagon-like peptide-1 protects beta-cells against apoptosis by increasing the activity of an IGF-2/IGF-1 receptor autocrine loop. Diabetes 58, 1816–1825 (2009).

61. Y. Li et al., Glucagon-like peptide-1 receptor signaling modulates beta cell apoptosis. J Biol Chem 278, 471–478 (2003).

62. N. B. Jorgensen et al., Exaggerated glucagon-like peptide 1 response is important for improved beta-cell function and glucose tolerance after Roux-en-Y gastric bypass in patients with type 2 diabetes. Diabetes 62, 3044–3052 (2013).

63. M. Salehi, R. L. Prigeon, D. A. D’Alessio, Gastric bypass surgery enhances glucagon-like peptide 1-stimulated postprandial insulin secretion in humans. Diabetes 60, 2308–2314 (2011).

64. J. Vidal, A. de Hollanda, A. Jimenez, GLP-1 is not the key mediator of the health benefits of metabolic surgery. Surg Obes Relat Dis 12, 1225–1229 (2016).

65. M. L. Vetter et al., GLP-1 plays a limited role in improved glycemia shortly after Roux-en-Y gastric bypass: a comparison with intensive lifestyle modification. Diabetes 64, 434–446 (2015).

66. J. Ye et al., GLP-1 receptor signaling is not required for reduced body weight after RYGB in rodents. Am J Physiol Regul Integr Comp Physiol 306, R352–362 (2014).

67. B. B. Boland et al., Combined loss of GLP-1R and Y2R does not alter progression of high-fat diet-induced obesity or response to RYGB surgery in mice. Mol Metab 25, 64–72 (2019).

68. J. D. Douros, J. Tong, D. A. D’Alessio, The Effects of Bariatric Surgery on Islet Function, Insulin Secretion, and Glucose Control. Endocr Rev 40, 1394–1423 (2019).

69. V. Poitout, R. P. Robertson, Glucolipotoxicity: fuel excess and beta-cell dysfunction. Endocr Rev 29, 351–366 (2008).

70. M. Y. Donath, S. E. Shoelson, Type 2 diabetes as an inflammatory disease. Nat Rev Immunol 11, 98–107 (2011).

71. J. Tian et al., Bile acid signaling and bariatric surgery. Liver Res 1, 208–213 (2017).

72. E. Akalestou, L. Genser, G. A. Rutter, Glucocorticoid Metabolism in Obesity and Following Weight Loss. Front Endocrinol (Lausanne) 11, 59 (2020).

73. J. Aron-Wisnewsky, J. Dore, K. Clement, The importance of the gut microbiota after bariatric surgery. Nat Rev Gastroenterol Hepatol 9, 590–598 (2012).

74. I. B. Leibiger, A. Caicedo, P. O. Berggren, Non-invasive in vivo imaging of pancreatic beta-cell function and survival - a perspective. Acta Physiol (Oxf) 204, 178–185 (2012).

75. M. A. Ravier, G. A. Rutter, Isolation and culture of mouse pancreatic islets for ex vivo imaging studies with trappable or recombinant fluorescent probes. Methods Mol Biol 633, 171–184 (2010).

